# MoTSE: an interpretable task similarity estimator for small molecular property prediction tasks

**DOI:** 10.1101/2021.01.13.426608

**Authors:** Han Li, Xinyi Zhao, Shuya Li, Fangping Wan, Dan Zhao, Jianyang Zeng

## Abstract

Understanding the molecular properties (e.g., physical, chemical or physiological characteristics and biological activities) of small molecules plays essential roles in biomedical researches. The accumulating amount of datasets has enabled the development of data-driven computational methods, especially the machine learning based methods, to address the molecular property prediction tasks. Due to the high cost of obtaining experimental labels, the datasets of individual tasks generally contain limited amount of data, which inspired the application of transfer learning to boost the performance of the molecular property prediction tasks. Our analyses revealed that simultaneously considering similar tasks, rather than randomly chosen ones, can significantly improve the performance of transfer learning in this field. To provide accurate estimation of task similarity, we proposed an effective and interpretable computational tool, named Molecular Tasks Similarity Estimator (MoTSE). By extracting task-related local and global knowledge from pretrained graph neural networks (GNNs), MoTSE projects individual tasks into a latent space and measures the distance between the embedded vectors to derive the task similarity estimation and thus enhance the molecular prediction results. We have validated that the task similarity estimated by MoTSE can serve as a useful guidance to design a more accurate transfer learning strategy for molecular property prediction. Experimental results showed that such a strategy greatly outperformed baseline methods including training from scratch and multitask learning. Moreover, MoTSE can provide interpretability for the estimated task similarity, through visualizing the important loci in the molecules attributed by the attribution method employed in MoTSE. In summary, MoTSE can provide an accurate method for estimating the molecular property task similarity for effective transfer learning, with good interpretability for the learned chemical or biological insights underlying the intrinsic principles of the task similarity.

## 1 Introduction

With the development of high-throughput experimental techniques in the fields of biology and chemistry [20], the number of available datasets of molecular properties has increased significantly over the past few years [22, 23, 13]. These data can provide comprehensive understanding into the molecular properties that are useful for drug design. In addition, the accumulating datasets can enable the design of accurate computational models for predicting the molecular properties and thus further facilitate and accelerate the drug discovery process. However, as huge experimental efforts are often required for obtaining large-scale molecular property labels, the available data of the majority of the desired properties are still extremely scarce. For example, although a preprocessed ChEMBL dataset [21, 7] contains 1310 bioassays covering over 400k small molecules, the data sizes of over 90% of the bioassays are below 1000. These insufficient datasets have limited the applications of data-driven computational models, especially deep learning models, into making accurate predictions of the molecular properties.

To alleviate the data scarcity problem, various strategies have been proposed to take advantage of the knowledge learned from related tasks, including transfer learning and multitask learning [29, 31]. Among these strategies, transfer learning is probably the most prevailing one, as its superior performance in the molecular property prediction tasks has already been relatively well validated [29, 31]. Given a task of interest (i.e., the target task), instead of training a model from scratch, transfer learning improves the prediction performance through finetuning a model pretrained from other related tasks (i.e., the source tasks). Nevertheless, the success of transfer learning is not always guaranteed, and a number of studies have indicated that transfer learning can also harm the prediction performance (termed negative transfer) [6, 26]. It has been observed that negative transfer usually occurs when the tasks are poorly related. That is, there exists only weak (or even no) similarity between source and target tasks [40]. Therefore, to facilitate the effective applications of transfer learning in predicting molecular properties and avoid the negative transfer problem, it is necessary to accurately measure the similarity between molecular property tasks to guide the source task selection and thus boost the prediction performance.

It is generally hard to explicitly or manually define the similarity between molecular property tasks, as fully understanding the behaviors of molecules in the chemical and biological systems is extremely difficult due to the high complexity of these systems. Fortunately, the data-driven computational methods enable us to define the task similarity in an implicit way. A widely-used method for measuring task similarity is ‘transferability’ [5, 39], which defines the similarity between two tasks as the amount of ‘advantage’ gained by using one task to improve the learning of another task. A successful example in the computer vision field is Taskonomy [39], which measures task transferability by performing transfer learning between each pair of tasks (i.e., one task acts as the source task, while the other task as the target task). The results have shown that incorporating task transferability can indeed improve the performance of transfer learning on the computer vision tasks. Moreover, as a fully automated method, Taskonomy derives the task similarity without involving any expert knowledge. In addition, the similarity tree constructed according to the calculated task transferability is highly consistent with human conception, indicating that task transferability captures the intrinsic relations between tasks in an implicit way.

Therefore, the task similarity can offer us a new perspective to understand the relations between tasks, and may even bring novel insights into certain chemical or biological tasks even their related mechanisms are not fully understood. This inspired us to develop a computational method for estimating the similarity between molecule property prediction tasks, which can not only guide the source task selection to avoid negative transfer and further improve the performance of transfer learning, but also help understand the relations between tasks and interpret the mechanisms behind them.

Despite the success of the transferability based methods, it is computationally expensive to estimate task transferability. It requires brute force search of the best source tasks through performing transfer learning between all possible pairs of tasks. Moreover, due to the black box nature of transfer learning, transferability provides only the implicit definition of task similarity with poor interpretability. To overcome these problems, we propose an interpretable computational molecular task similarity estimator, MoTSE, to efficiently measure the task similarity, based on the assumption that two tasks should be similar if the knowledge learned by their task-specific models is close to each other. The main idea of MoTSE is to project individual tasks into vectors in a latent space, and then use the distances between the vectors to derive the similarity between the tasks. The projection is realized by embedding the task-related local and global knowledge learned by the pretrained GNN models into the vectors, with the help of a set of unlabeled probe data.

In our paper, extensive computational evaluation demonstrated that the task similarity estimated by MoTSE successfully can guide the source task selection in transfer learning, with superior performance over several baseline methods including multitask learning and training from scratch, under several quantum mechanical and biophysical property datasets. The generalization ability of MoTSE was further validated using the learned task similarity from the pretrained GNN models to boost performance of other models, and applying the task similarity estimated using one quantum mechanical dataset to another dataset containing the same tasks but with a different data distribution. In addition, comprehensive computational experiments demonstrated that MoTSE is more applicable in practice, especially for large task sets, with less time and resource required to obtain reliable results. Finally, we analyzed the interpretability of the estimated similarity between tasks and demonstrated the ability of our model to bring biological or chemical insights into understanding the molecular property prediction tasks.

## 2 Method

### 2.1 Notation and Problem Setting

Suppose that we are given a set of tasks 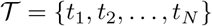, where *N* stands for the total number of tasks involved. Accordingly, we have a set of datasets 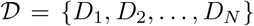, where 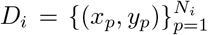 stands for the dataset related to task *t_i_*, and (*x_p_, y_p_*) represents the *p*-th pair of molecule and its label for task *t_i_* and *N_i_* stands for the size of *D_i_*. We use a graph representation to encode the features of a molecule. More specifically, given a molecule, we use RDKit (http://www.rdkit.org) to extract its graph representation 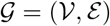, where 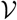 stands for the set of nodes (i.e., heavy atoms) and ***ɛ*** stands for the set of edges (i.e., covalent bonds). We mainly represent individual atoms by their one-hot encodings of atoms types. In particular, we use 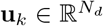 to represent the *k*-th node in 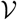, where *N_d_* stands for the dimension of node features (i.e., the number of atom types). Our goal mainly lies in the following two folds: (1) efficiently calculate the similarity between each pair of tasks in 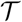 without involving any expert knowledge; and (2) given a target task 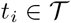 of interest, use the derived similarity to recommend the task *t_j_* that is the most similar to task *t_i_* as the source task so that the prediction performance of transfer learning on task *t_i_* can be optimized.

### 2.2 The MoTSE Framework

Given a task set 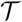 and the corresponding datasets 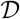 for individual tasks, we introduce MoTSE, an efficient computational method to estimate the similarity between molecular property prediction tasks. The core idea of our method is to embed the task-related knowledge extracted from the pretrained GNNs of each task into a unified task space and then calculate the similarity by measuring the distance between the embedded vectors of the corresponding tasks. The overall workflow of our proposed method is illustrated in Figure 1, and the key steps will be described in detail below.

**Fig. 1.**
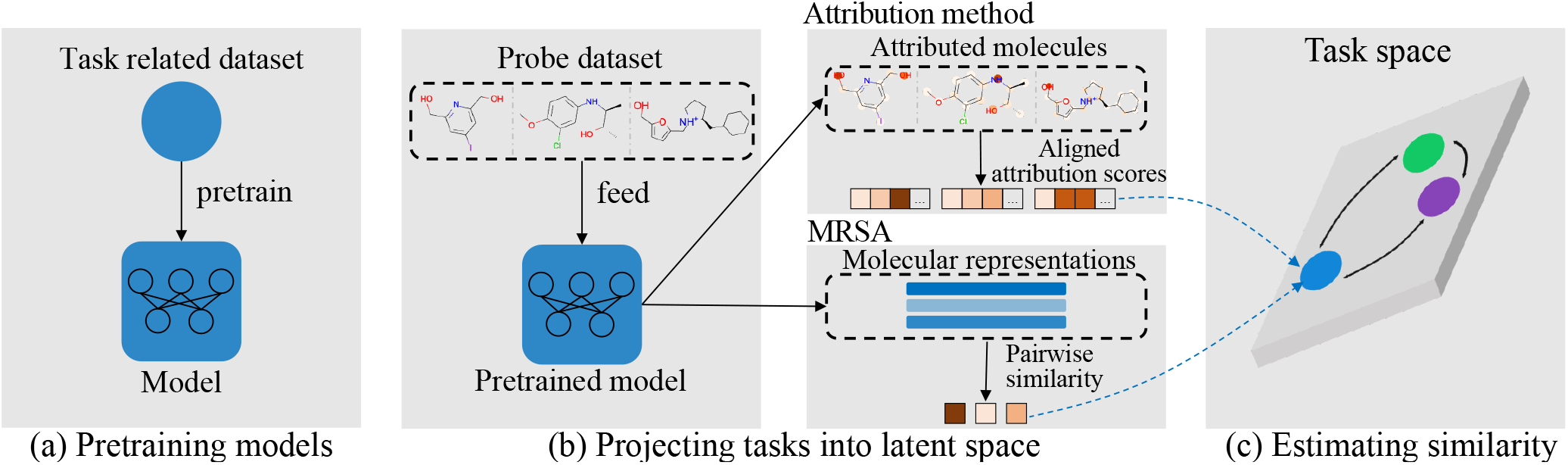
An illustrative diagram of MoTSE. (a) MoTSE first pretrains a GNN for each task using the corresponding dataset. (b) MoTSE then projects the task-related knowledge extracted from the pretrained GNNs of individual tasks into a task space with a probe dataset (a set of randomly selected unlabeled molecules). The knowledge extraction is achieved by two methods: an attribution method extracting the task-related local knowledge by assigning importance scores to individual atoms in a molecule; and a molecular representation similarity analysis (MRSA) method extracting the task-related global knowledge by pair-wisely measuring the similarity between molecular representations. (c) Finally, MoTSE calculates the similarity between tasks by measuring the distance between the embedded vectors of the corresponding tasks in the embedded task space.

### 2.3 Key Steps

#### Step 1: Pretraining the Task-Specific GNNs

The calculation of the similarity between tasks can be regarded as measuring the intrinsic knowledge that need to be learned from these tasks. Since deep learning models, especially the GNNs, have shown their superior capability of extracting molecular representations and modeling various kinds of molecular properties [8, 19, 38], we adopt a data-driven approach to capture the hidden knowledge contained in individual tasks. More specifically, for each task 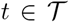, we first pretrain a GNN model *m* = *p*(*e*(·)) from its corresponding dataset *D*, where *e*(·) acts as an encoder to extract the latent feature representations of the molecule graphs and *p*(·) serves as a classifier or regressor (implemented through a fully connected neural network) to make prediction for *t*. After pretraining, we consider that the knowledge of each task is included in the corresponding trained GNN.

#### Step 2: Projecting Tasks into Task Space

After pretraining task-specific GNNs for individual tasks, the problem of measuring the similarity between two tasks is converted into finding a way to represent and measure the knowledge enclosed in the pretrained GNNs. To this end, MoTSE projects each task into a task space (i.e., as a vector representation) based on the task-related knowledge extracted from the pretrained models. More specifically, we first define a probe dataset 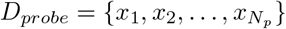, which is a set of randomly selected unlabeled molecules from a small molecule library, where *N_p_* denotes the number of molecules contained in *D_probe_*. This probe dataset is shared across all tasks involved and then acts as a proxy in the process of projecting the knowledge of individual tasks into a same task space.

Here, we present two projection methods, including an attribution method and a molecular representation similarity analysis (MRSA) method, to extract the vector representations of the task-specific knowledge from the pretrained GNNs using the probe dataset.

##### Attribution Method

The main idea of an attribution method is to assign importance scores for individual input features. For a specific molecular property prediction task, we first use this attribution method to assign importance scores for individual atoms in a molecule and detect which functional groups in a molecule are important for the learning task. Here, the specific attribution method we use is Gradient*Input [30], which refers to a first-order Taylor approximation of how the output will change if a specific input feature is set to zero, thus indicating the importance of this input feature with respect to the output.

More specifically, given a graph representation 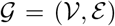 of molecule *x* ∈ *D_probe_*, the attribution score *a_k_* of the feature vector of the *k*-th atom **u**_*k*_ with respect to the task *t* can be computed as:

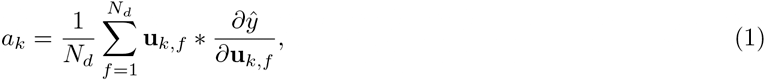

where **u**_*k,f*_ is the *f*-th element of the feature vector **u**_*k*_, *N_d_* is the dimension of the input atom features, and 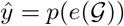 is the prediction result of *x* for task *t*.

Here, the attribution score *a_k_* of the *k*-th atom is derived based on the mean of the attribution scores of all dimensions of the atom features. After assigning the attribution scores to individual atoms of molecule *x*, we obtain an attribution vector denoted by 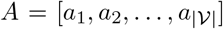, where 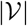 stands for the number of atoms in *x*. Similarly, we can derive the attribution vectors of all molecules in the probe dataset 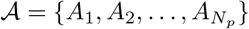, by applying the above attribution method to every molecule in *D_probe_*.

##### Molecular Representation Similarity Analysis

As the attribution method scores each atom separately without considering the global information of molecules (e.g., the knowledge contained in the latent feature representation of the whole molecule), we define the knowledge extracted by the above attribution method as local knowledge. Here, we present a molecular representation similarity analysis (MRSA) [4, 9] as the second method to extract global knowledge from the pretrained GNNs using the probe dataset *D_probe_*. In particular, we compute the pairwise correlation between the hidden molecule representations generated by the pretrained GNNs (i.e., the outputs of the encoders of the pretrained GNNs). We define such knowledge extracted by the MRSA method as global knowledge, as the latent molecular representations contain the indispensable information of the whole molecules with respect to the downstream task and thus the pairwise correlation between them can be effectively used to represent their discrepancy in the latent molecular representation space.

More specifically, for task *t* and the encoder *e* from the corresponding model, we first perform forward propagation for all molecules in *D_probe_* to generate their latent molecular representations 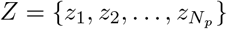, where *z_m_* stands for the latent molecular representation of molecule *x_m_* ∈ *D_probe_*. Then for each pair of molecular representations *z_m_* and *z_n_* (*m* ≠ *n*), we compute their correlation score *r_m,n_*, that is,

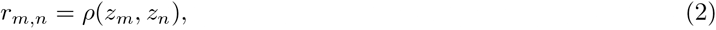

where *ρ* stands for the Pearson’s correlation coefficient. After that, we obtain a molecular representation correlation vector 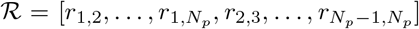 as another knowledge representation of task *t*, where *N_p_* stands for the size of the probe dataset.

For individual tasks in 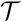, MoTSE adopts the above two knowledge extraction methods to extract both local and global task-related knowledge and then projects them as vectors into two latent task spaces 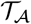 and 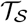 constructed by the attribution method and the MRSA approach, respectively.

#### Step 3: Estimating Task Similarity

Once step 2 is completed, for each pair of tasks 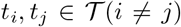, their similarity can be computed based on the latent task spaces 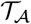 and 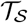:

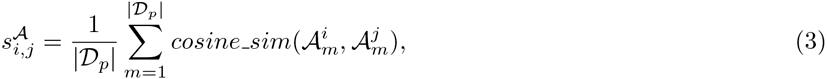

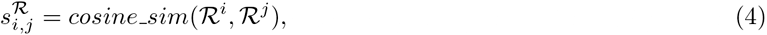

where 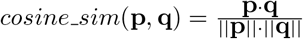 stands for the cosine similarity between two vectors.

These two kinds of task similarity are calculated under different assumptions. As the attribution method mainly aims to extract local knowledge, the assumption behind 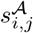 is that similar tasks should have similar attribution scores for individual atoms in a molecule, which is consistent with the perception of biologists/chemists. On the other hand, MRSA mainly aims to extract the global knowledge, and 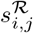 measures the similarity on the basis that similar tasks should result in similar latent molecular representation spaces, which are determined based on the knowledge learned by GNNs. To fully exploit the merits of both similarity estimation methods, we unify them into a more comprehensive one:

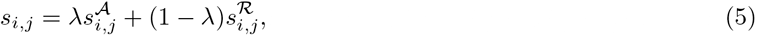

where λ stands for the weighting factor between two types of similarity.

## 3 Results

### 3.1 Experimental Settings

#### Datasets

We used two representative datasets QM9 [23] and PCBA [13, 24] to evaluate the effectiveness of our proposed method. QM9 is a dataset that provides quantum mechanics properties, such as geometric, energetic, electronic and thermodynamic properties of roughly 130K small molecules with up to nine heavy atoms, associating with 12 regression tasks [23]. PCBA is a dataset consisting of biological activities of small molecules generated by high-throughput screening, associating with 128 classification tasks [24]. The transfer learning strategy adopted in this work included a pretraining stage on the source tasks and a finetuning stage on the target tasks. To demonstrate the effectiveness of incorporating the task similarity in boosting the prediction performance of transfer learning, we further preprocessed these two datasets to (1) mimic a scenario of transfer learning, in which the data size of the source task was relatively larger than that of the target task, and (2) reduce the influence of other factors (e.g., data size and label imbalance) that might affect the performance of transfer learning and thus only focus on the effect of task similarity [40]. In particular, we first created two subsets QM9_10*k*_ and PCBA_10*k*_ as the datasets for the source tasks. QM9_10*k*_ was processed by randomly sampling 10k data for all 12 tasks from QM9. As PCBA is an extremely unbalanced dataset, we created PCBA_10*k*_ by first selecting the 23 tasks that contained more than 5k positive and 5k negative labels and then constructed a balanced 10k dataset by randomly sampling 5k positive and 5k negative samples for each of these tasks. QM9_10*k*_ and PCBA_10*k*_ were further partitioned into training (8k), validation (1k) and test (1k) sets. Then we created datasets QM9_*n*_ and PCBA_*n*_ as the datasets for the target tasks (i.e., tasks used in finetuning), where *n*<10*k* denotes the sizes of the datasets. We varied *n*1*k,* 2*k,* 3*k,* 4*k,* 5*k* to evaluate the prediction performance of transfer learning under different data sizes of target tasks. To avoid data leakage during transfer learning, the training and validation sets of QM9_*n*_ (PCBA_*n*_) were obtained by randomly sampling 0.8*n* and 0.1*n* samples from the training and validation sets of QM9_10*k*_ (PCBA_10*k*_), respectively. The test set of QM9_*n*_ (PCBA_*n*_) was the same as that of QM9_10*k*_ (PCBA_10*k*_) for a fair comparison.

#### Metrics

We mainly used R^2^ (the coefficient of determination) and AUROC (the area under receiver operating characteristic) to evaluate the prediction performance of models for regression and classification tasks, respectively. R^2^ measures how well the predictions fit the true labels, while AUROC evaluates the aggregation classification performance across all possible thresholds. To further evaluate whether MoTSE can recommend the correct source tasks, we also introduce *P*@*K* and *R*@*K*, which have been widely used as effective metrics for the evaluation of recommendation systems. *P*@*K* stands for the proportion of recommended items in the top K set that are correct, and *R*@*K* represents the proportion of correct items found in the top K recommendations.

#### Pretraining and Finetuning

For each task in QM9_10*k*_ and PCBA_10*k*_, we pretrained a GNN for finetuning and task similarity estimation. To ensure that no bias was introduced, we preserved the same model architecture and training details across all the tasks. More specifically, we adopted the graph convolutional network (GCN) [15] implemented by DGL [35] as the encoder and a three-layer perceptron as the predictor. Adam optimizer [14] was employed for optimizing the model parameters by gradient descent. During the finetuning stage, all the model parameters and the training details were the same as in pretraining. The training details are provided in Supplementary Section 1.1.

#### Probe Dataset

In our computational experiments, we also constructed a probe dataset by randomly sampling 500 small molecules from a Zinc dataset preprocessed by previous research [32, 17] for task similarity estimation. This probe dataset was shared across all formerly defined tasks and acted as a proxy in the process of projecting individual tasks into the latent task space as described in Section 2. More details about the effects of the randomness and the size of probe dataset on the performance of MoTSE are provided in Supplementary Sections 2.2 and 2.5.

#### Task Similarity Estimation

For each of the formerly defined datasets, we estimated the similarity between individual pairs of tasks using MoTSE as described in Section 2. We set the weighting factor *λ* to 0.8 in the following computational experiments. We denoted the similarity estimated using QM9_10*k*_ and PCBA_10*k*_ as MoTSE_10*k*_ and the similarity estimated using QM9_*n*_ and PCBA_*n*_ as MoTSE_*n*_, where *n* ∈ {1*k,* 2*k,* 3*k,* 4*k,* 5*k*}. To comprehensively evaluate the effect of incorporating task similarity on the performance of transfer learning, we also introduced another kind of task similarity measurement, that is, task transferability. More specifically, given two tasks 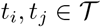 (*i* ≠ *j*), the task transferability from *t_i_* to *t_j_* can be defined as the prediction performance (i.e., R^2^ or AUROC) of transfer learning when using *t_i_* as the source task and *t_j_* as the target task. Then we defined transferability_*n*_ as the task transferability estimated when the tasks in QM9_10*k*_ (PCBA_10*k*_) acted as source tasks and the tasks in QM9_*n*_ (PCBA_*n*_) acted as target tasks, where *n* ∈ {1*k,* 2*k,* 3*k,* 4*k,* 5*k*}.

### 3.2 Performance Evaluation

In this section, we first demonstrated that incorporating task similarity information (i.e., the task similarity estimated by MoTSE or task transferability) is crucial for molecular property prediction in the transfer learning process. Given a task set 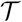 and task similarity between individual pairs of tasks in 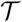, for a target task 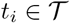, task similarity information can be used to improve the transfer learning performance on *t_i_* by selecting a source task *t_j_* (*i* ≠ *j*) with the highest similarity to *t_i_*. To demonstrate this point, we evaluated the prediction performance of transfer learning with the guidance of such a source task selection process on QM9 and PCBA datasets. More specifically, we used the formerly defined QM9_10*k*_ and PCBA_10*k*_ as the datasets for pretraining and QM9_*n*_ and PCBA_*n*_ (*n* ∈ 1*k,* 2*k,* 3*k,* 4*k,* 5*k*) as the datasets for finetuning, to evaluate the prediction performance of transfer learning under different task similarity estimation methods. Then for each target task in datasets QM9_*n*_ and PCBA_*n*_ (*n* ∈ {1*k,* 2*k,* 3*k,* 4*k,* 5*k*}) for finetuning, we carried out the following four settings for source task selection:

- MoTSE_10*k*_-3: Selecting source tasks using the task similarity computed by MoTSE_10*k*_. For each target task, we selected top three tasks with the highest similarity computed by MoTSE_10*k*_. Next, we used each of them as the source task for transfer learning, and then took the best finetuning result as the final one.
- MoTSE_*n*_-3: Selecting source tasks using the task similarity computed by MoTSE_*n*_, where *n* denotes the size of current dataset for finetuning, and then performing the same source task selection and finetuning processes as in MoTSE_10*k*_-3.
- Best-finetune: Selecting source tasks using transferability_*n*_, where *n* denotes the size of current dataset for finetuning. As defined in Section 3.1, given a target task, the source task leading to higher prediction performance has higher task transferability with the target task. Therefore, if we select the source task with the highest transferability, we can always obtain the best transfer learning performance, which can serve as an upper bound of the prediction performance of transfer learning with different source tasks selection schemes.
- Random-3: Selecting source tasks randomly. For each target task, we randomly selected three source tasks and performed the same finetuning processes as in MoTSE_10*k*_-3 and MoTSE_*n*_-3. To eliminate the influence of the randomness on the results, we reported the expected values as the final results by taking the average of all possible results.

For comparison, we also introduced the following two baselines:

- Scratch: training the target task from scratch (i.e.,without incorporating information from other tasks).
- Multitask: training the target task and all source tasks in a multitask learning fashion.

For both QM9 and PCBA, we plotted the R^2^ and AUROC of the above settings under different data sizes of target tasks (Figure 2a-b). Our test results showed that all the transfer learning settings outperformed the training scheme from the scratch setting by a large margin, indicating that transfer learning was indeed a powerful strategy for improving prediction performance of tasks with limited data. Also, selecting source tasks according to task similarity (i.e., Best-finetune, MoTSE_10*k*_-3 and MoTSE_*n*_-3) achieved much better prediction performance than randomly selecting the source tasks (i.e., Random-3). In addition, the perfect transfer learning setting (i.e, Best-finetune) outperformed multitask learning, which suggested that incorporating a relatively larger number of tasks without assessing their similarity with the target tasks in the training process (i.e., multitask learning) may not necessarily achieve better results and even impede the prediction performance. In other words, the performance of multitask learning might be worse than that of training from scratch (Figure 2a). According to these observations, we can draw the conclusion that transfer learning with task similarity provides an effective strategy for improving molecular property predictions.

**Fig. 2.**
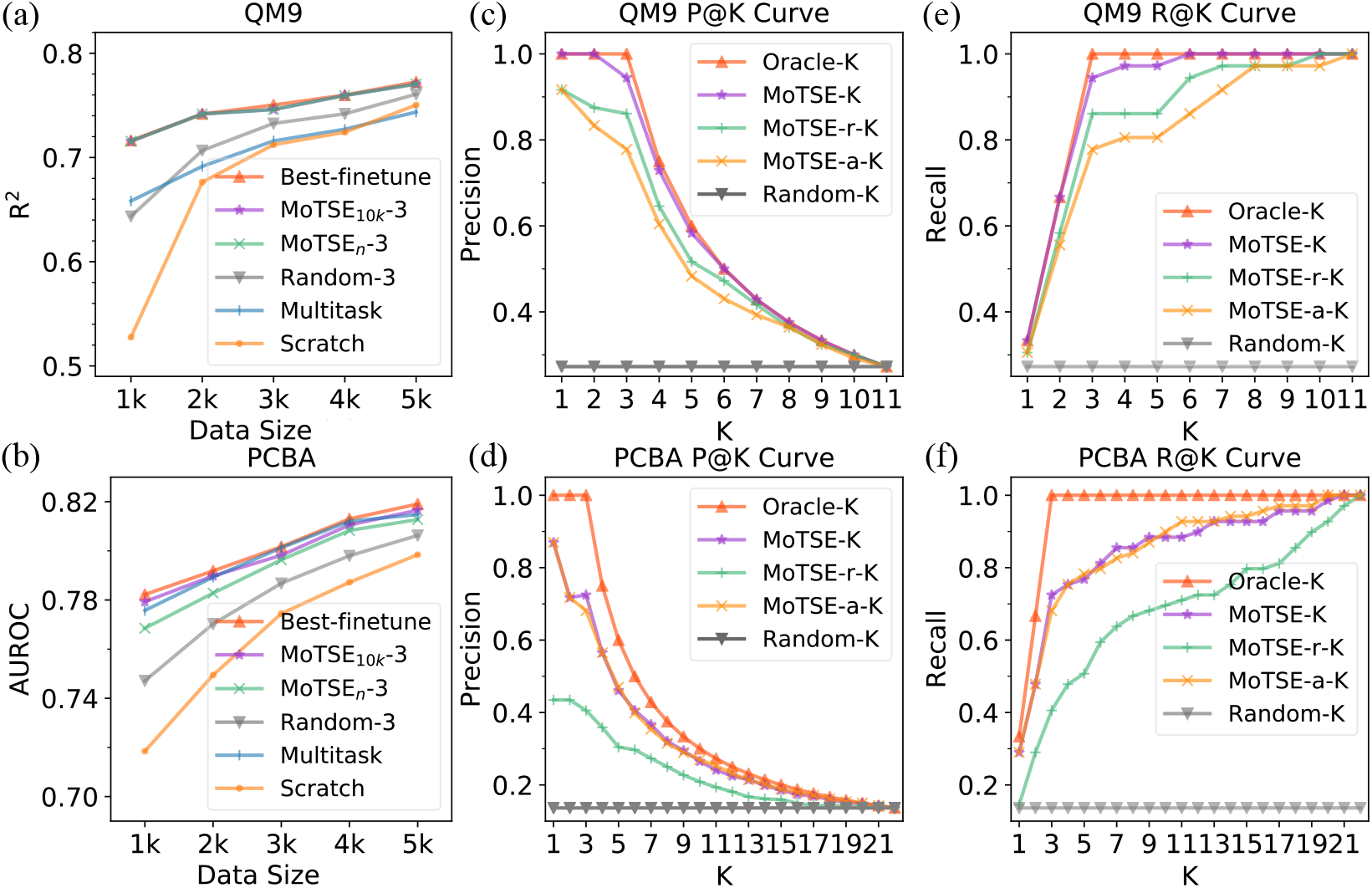
Performance of transfer learning with task similarity estimated by MoTSE and other strategies. (a,b) The prediction performance on QM9 (a) and PCBA (b) of transfer learning with four source different task selection schemes (i.e., Best-finetune, MoTSE_10*k*_-3, MoTSE_*n*_-3 and Random-3) and two baselines (i.e., Scratch and Multitask). (c-f) The recommendation performance of the similarity estimated by MoTSE on QM9 and PCBA datasets. (c,e) The *P*@*K* and *R*@*K* curves evaluated on QM9. (d,f) The *P*@*K* and *R*@*K* curves evaluated on PCBA. MoTSE-r and MoTSE-a stand for two variants of MoTSE, in which MoTSE-a estimated the task similarity only using the attribution method (i.e., Eq. 3), and MoTSE-r estimated the task similarity only using MRSA (i.e., Eq. 4). Random recommendation was introduced as a baseline denoted as Random, and the ideal method denoted as Oracle, which always recommends the correct tasks (i.e., recommending tasks according to transferability_1*k*_).

Moreover, we obtained close results when performing the source task selection using the similarity estimated by MoTSE (i.e., MoTSE_10*k*_-3) and task transferability (i.e., Best-finetune), even when the MoTSE similarity was estimated using only limited data (i.e., MoTSE_*n*_-3). This result indicated that the similarity estimated by MoTSE provided a nearly perfect guidance for the source task selection process. Note that, by increasing the number of tasks chosen in the source task selection process, we can gradually improve the prediction performance of transfer learning with the guidance of similarity estimated by MoTSE until the best performance (i.e., Best-finetune) was reached (Figure S2 and Figure S3).

To further demonstrate the effectiveness of the similarity estimated by MoTSE in the source task selection process, we considered MoTSE as a recommendation system (i.e., recommending a source task for a given target task), where each target task acted as a query, and the tasks with the top three highest transferability_1*k*_ were considered to be the answers to the query. Then we adopted *P*@*K* and *R*@*K* to evaluate whether MoTSE can accurately recommend tasks with high transferability. In addition, to demonstrate the importance of each module of MoTSE, we also introduced two variants of our method, including MoTSE-a which only used the attribution method when estimating the task similarity (i.e., Eq. 3) and MoTSE-r which only used MRSA when estimating the task similarity (i.e., Eq. 4).

The *P*@*K* and *R*@*K* curves showed that the source tasks with the top three highest transferability can be accurately recommended by the similarity estimated by MoTSE (Figure 2c-f). Detailed analyses revealed that among 11 out of 12 tasks in QM9 and 21 out of 23 tasks in PCBA, the source tasks with the highest transferability can be included in the top three recommendations from MoTSE, which explained why MoTSE achieved comparable prediction performance with Best-finetuning on QM9 and PCBA. Moreover, MoTSE outperformed MoTSE-a and MoTSE-r by a considerable margin, which implied the necessity of using both the local and global knowledge for measuring the task similarity. We also observed that the performance of MoTSE-r dramatically dropped on the PCBA dataset. This may be explained by the fact that the bioacitivity properties in PCBA are mostly determined by the local features (e.g., functional groups), while MoTSE-r mainly focused on the global information.

MoTSE is much more efficient than the estimation of task transferability and thus more applicable in practice. Task transferability is generally much more expensive to calculate, especially when the number of tasks in 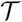 and their sizes are large, because we have to perform transfer learning for each pair of tasks in this setting. However, MoTSE runs much faster as it only pretrains a model for each task and conducts one time forward-and-backward propagation for each model on a relatively small probe dataset. This improvement is significant since it is a common scenario in which 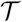 the task set is very large. In our computational experiments, after pretraining, MoTSE can estimate pairwise similarity within 5 minutes between tasks on PCBA using CPU, while measuring transferability required about 20 GPU hours with one GTX 1080 Ti GPU. More details about the computational complexity analysis of MoTSE and task transferability estimation can be found in the Supplementary Section 1.4.

The above results demonstrated that incorporating task similarity (i.e., task transferability and the similarity estimated by MoTSE) is indeed of great significance for boosting the performance of molecular property prediction, and MoTSE is an effective and a more practically applicable task similarity estimation method, especially for the applications in large task sets.

### 3.3 Generalizability of MoTSE

We previously have shown that the similarity estimated by MoTSE on GCN can effectively guide the transfer learning of a GCN and thus boost the prediction performance on the molecular property prediction tasks on QM9 and PCBA. In this section, we further demonstrated that the similarity estimated by MoTSE using a specific model (e.g., GCN) or specific dataset (e.g., QM9) is generalizable to guide the transfer learning process on other models or datasets with different data distributions and thus can yield better prediction performance.

To comprehensively evaluate the generalizability of the similarity estimated by MoTSE across different models, we employed three models, including another graph-based model, i.e., graph attention network (GAT) [34], a ECFP (extended connectivity fingerprints) [25] based fully-connected network (FCN) and a SMILES (simplified molecular input line entry specification) [36] based recurrent neural network (RNN) (more details of these three types of models can be found in Supplementary Section 1.1). With the guidance of the similarity estimated by MoTSE using a GCN (i.e., MoTSE_10*k*_-3 and MoTSE_*n*_-3 as defined in Section 3.1), we evaluated the performance of transfer learning of these three types of models on the QM9 and PCBA datasets. In particular, we first pretrained models on QM9_10*k*_ and PCBA_10*k*_, and then finetuned models on QM9_*n*_ and PCBA_*n*_, where *n* 1*k,* 2*k,* 3*k,* 4*k,* 5*k*. Next, for individual models, we plotted the data sizes versus R^2^ and AUROC on both QM9 (Figure 3a-c) and PCBA (Figure 3e-g).

**Fig. 3.**
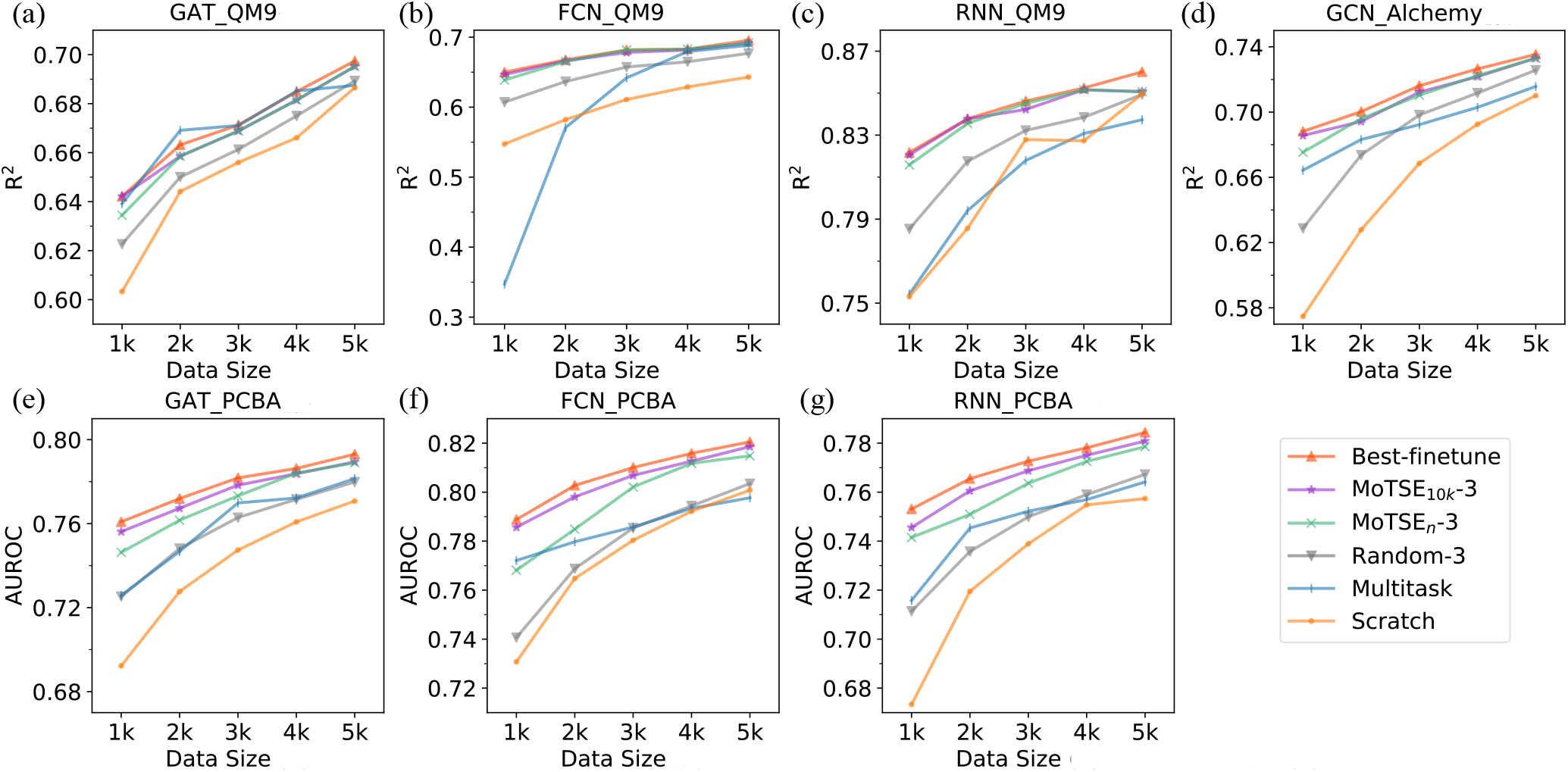
Generalizability of the task similarity estimated by MoTSE. The task similarity computed by MoTSE_10*k*_ and MoTSE_*n*_ as defined in Section 3.1 were used to guide the transfer learning process on different types of models or datasets with distinct data distributions. (a-c) The prediction performance of graph attention network (GAT) (a), fully-connected network (FCN) (b) and recurrent neural network (RNN) (c) on the QM9 dataset under transfer learning with the guidance of the similarity estimated by MoTSE using a GCN model. (d) The prediction performance of a GCN on Alchemy under transfer learning with the guidance of the similarity estimated by MoTSE using the QM9 dataset. (e-g) The prediction performance of GAT (e), FCN (f) and RNN (g) on the PCBA dataset under transfer learning with the guidance of the similarity estimated by MoTSE using a GCN model. For comparison, we also provided the results of Best-finetune, Random-3, Scratch and Multitask as defined in Section 3.2 for individual tests.

In addition, to evaluate the generalizability of the similarity estimated by MoTSE across datasets with different data distributions, we employed Alchemy [2], which has the same tasks but shares different data distribution with QM9, that is, QM9 contains molecules comprising up to 9 non-hydrogen atoms while the molecules in the Alchemy dataset consist of non-hydrogen atoms ranging from 9 to 14. We preprocessed Alchemy and created Alchemy_10*k*_ and Alchemy_*n*_ using the same manner as in QM9_10*k*_ and QM9_*n*_, where *n* ∈ {1*k,* 2*k,* 3*k,* 4*k,* 5*k*}. With the guidance of the similarity estimated by MoTSE using QM9 (i.e., MoTSE_10*k*_-3 and MoTSE_*n*_-3 as defined in Section 3.1), we evaluated the performance of transfer learning of a GCN on Alchemy. We first pretrained models on Alchemy_10*k*_, and then finetuned models on Alchemy_*n*_, where *n* ∈ {1*k,* 2*k,* 3*k,* 4*k,* 5*k*}. After that we plotted the data sizes versus R^2^ on the Alchemy dataset (Figure 3d).

For comparison, we also provided the results of Best-finetune, Random-3, Scratch and Multitask as defined in Section 3.2 for individual tests (Figure 3). The results indicated that by incorporating task similarity, transfer learning (i.e., Best-finetune, MoTSE_10*k*_-3 and MoTSE_*n*_-3) still outperformed Random-3, Scratch and Multitask on different models and datasets, which further verified the significance of the task similarity in the prediction of molecular properties. Moreover, the similarity estimated using the GCN model can effectually guide the transfer learning on other types of models, even if they are not graph-based (i.e., ECFP and SMILES based) models (Figure 3a-c and Figure 3e-g). In addition, the similarity estimated from the QM9 dataset can also effectually guide the transfer learning on the Alchemy dataset (Figure 3d). These results demonstrated the generalizability of the similarity estimated by MoTSE across different types of models and datasets with distinct data distributions, indicating that MoTSE can capture the intrinsic similarity between tasks without relying on the models used or datasets tested.

## 4 Interpretability of MoTSE

The above results have shown that MoTSE performed well on estimating task similarity and it is also of great necessity to explore how MoTSE interprets the estimated task similarity. Unlike transferability which works as a black box, the attribution method (Section 2.3) employed in MoTSE can assign an importance score for each atom in a molecule to the final prediction (i.e., attributed molecule). Through visualizing the attributed molecules, we can open the box and thus provide useful hints into understanding the task-related knowledge extracted by MoTSE.

### 4.1 Interpreting Similarity between Physical Chemistry Tasks

We first explored how MoTSE interprets the similarity between four well-studied physical chemistry tasks, including NHA (number of hydrogen acceptors contained in a molecule), NHD (number of hydrogen donors contained in a molecule), NOCount (number of nitrogen (N) and oxygen (O) atoms contained in a molecule), and NHOHCount (number of N and O atoms that are covalently bonded with hydrogens in a molecule). From the chemical perspective, NHA is more similar to NOCount than NHOHCount, and NHD is more similar to NHOHCount than NOCount, as those N and O atoms without any covalently bonded hydrogens can serve as only hydrogen bond acceptors but not donors.

We first constructed a dataset containing 10k molecules labeled with the above four properties. More specifically, the molecules in this dataset were randomly sampled from the Zinc dataset preprocessed by previous research [32, 17] and all the properties were computed by RDKit (http://www.rdkit.org). Then we estimated the similarity between these tasks using MoTSE and looked into the attributed molecules. The similarity between tasks derived by MoTSE (Figure 4) was completely in accordance with the existing prior knowledge. More specifically, NHA had the highest similarity with NOCount and NHD had the highest similarity with NHOHCount. More importantly, MoTSE precisely assigned high attribution scores to the target atoms (e.g., N and O for NOCount), and similar tasks tended to assign similar attribution scores to the same atoms in molecules (Figure 4). These observations interpreted how MoTSE estimated similarity between different tasks and indicated that our method indeed can capture the chemical concepts behind the tasks.

**Fig. 4.**
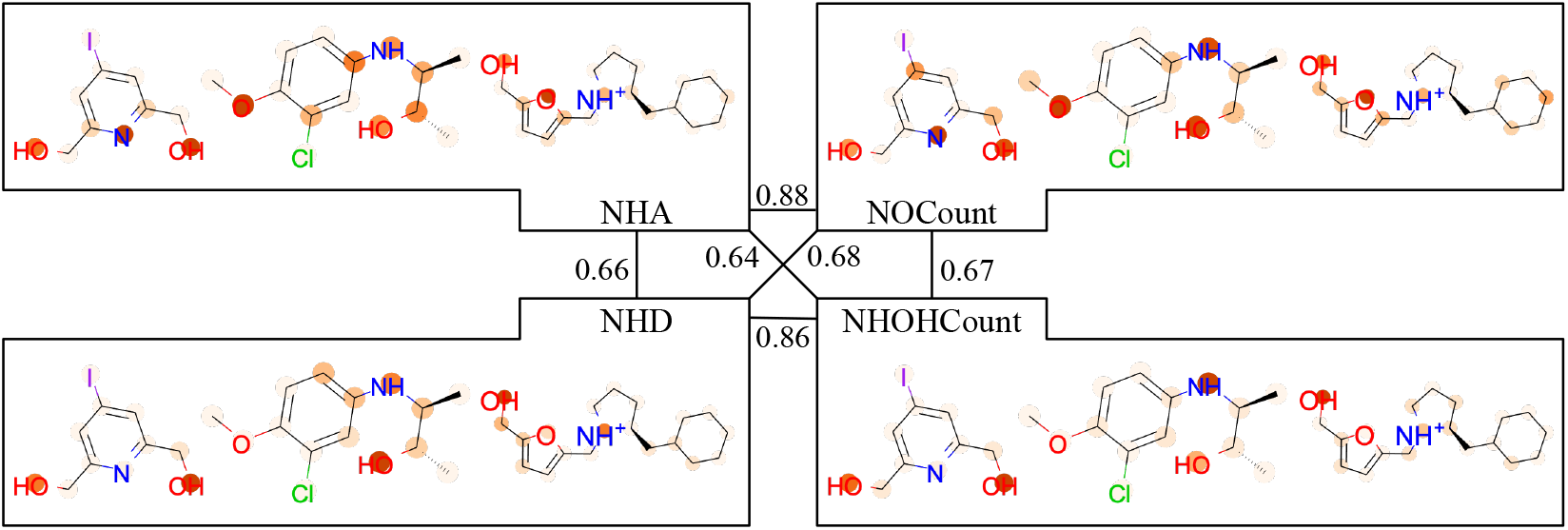
The similarity between four physical chemistry tasks (see the main text for their definitions) and the visualization of three example molecules attributed by our attribution method. The numbers between tasks denote the task similarity derived from MoTSE. In the attributed molecules, darker color represents higher importance.

### 4.2 Interpreting Similarity between the Tasks of Estimating the Bioactivities of Compounds against Cytochrome P450 Isozymes

To further demonstrate the interpretability of MoTSE, we carried out a more challenging experiment, which included another five tasks of predicting the bioactivities of small molecules against cytochrome P450 isozymes. The cytochrome P450 (CYP) family plays important rules in drug metabolism, especially for five isozymes - 1A2, 2C9, 2C19, 2D6 and 3A4 [3, 37]. Here we obtained the binary bioactivity labels between around 17k molecules and the above five CYP isozymes from a preprocessed ChEMBL dataset [21, 7, 33].

According to the similarity estimated by MoTSE, we first constructed a similarity tree using the hierarchical agglomerative clustering algorithm [10, 11] (Figure 5a). We found that the structure of this tree was exactly the same as that derived by self organizing maps (SOMs) [16, 27, 28] offered in a previous research [33], in which SOMs of individual isozymes were constructed based on the structural similarity of compounds and reflected the activity patterns (i.e., the scaffolds enriched in active and inactive compounds) of corresponding isozymes. According to this observation, we expected that MoTSE may capture the activity patterns of the CYP bioactivity prediction tasks and fully exploit such knowledge to estimate the similarity between these tasks. To validate this hypothesis, we further visualized several examples of attributed molecules. As shown in Figure 5b-f, we found that similar tasks tended to share the same active patterns (e.g., CYP 2C9 and CYP 2C19 shared the same five active patterns). In addition, the active patterns highlighted by our attribution method can be supported by previous researches [12, 18, 33]. For example, the substructure in Figure 5b was also previously considered as an active pattern for CYP 2C9, CYP 2C19 and CYP 3A4 by substructure searching [33] and fingerprint analysis [18]. Moreover, the similarity between CYP2C9 and CYP2C19 estimated by MoTSE was the highest among all pairs of CYP isozymes, which can also be supported by the fact that CYP 2C9 and 2C19 geneically share the most (91%) sequence homology [1]. The above results demonstrated that MoTSE can successfully extract task-related knowledge and thus accurately estimate the intrinsic similarity between tasks (e.g., the similarity between active patterns of compounds and the genetic similarity between CYP isozymes). Therefore, MoTSE can provide an insightful perspective to help understand the mechanisms underlying the molecular properties.

**Fig. 5.**
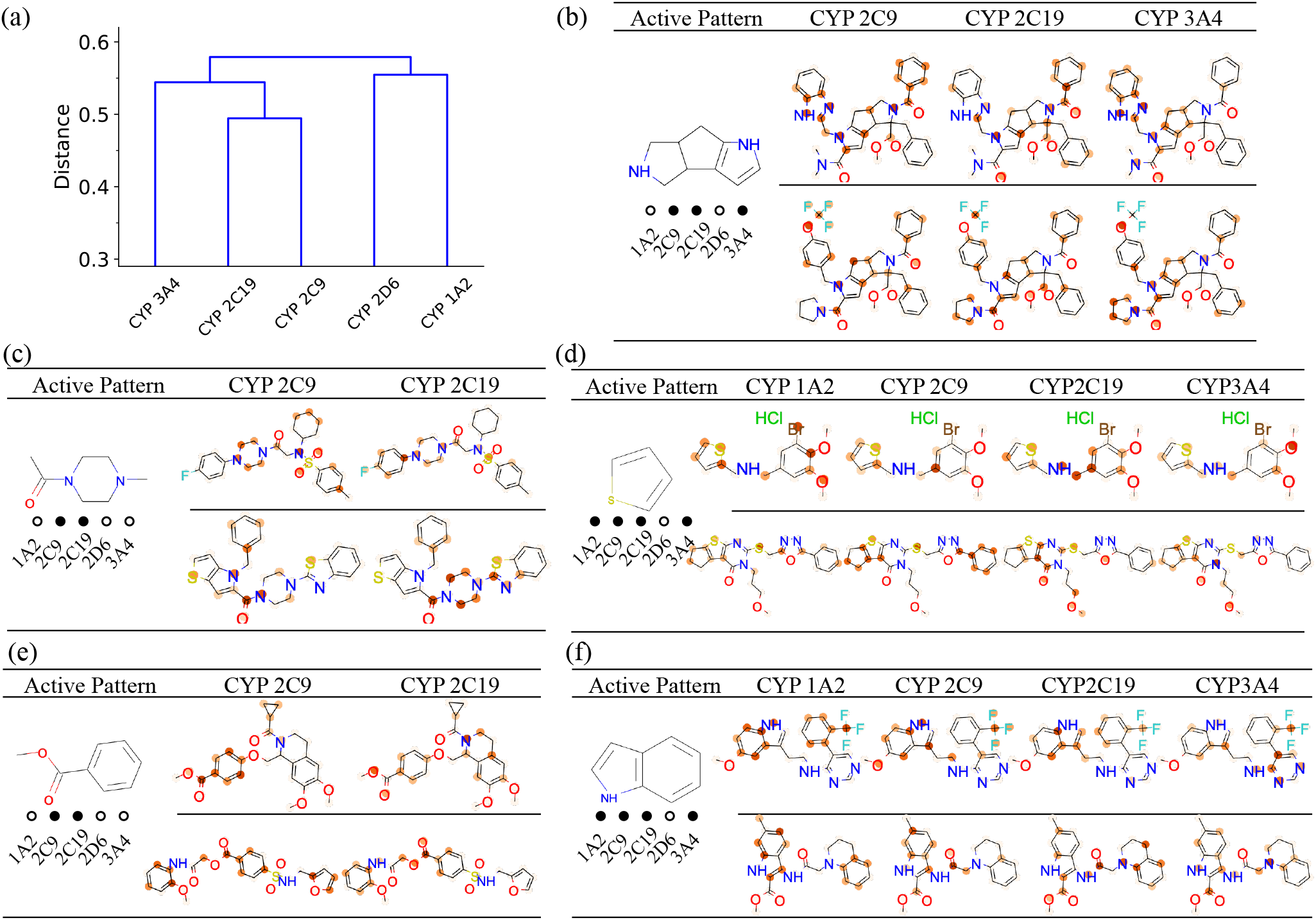
Interpreting the task similarity of MoTSE in predicting the bioactivities of compounds against CYP isozymes. (a) Task similarity tree constructed using the similarity estimated by MoTSE. (b-f) Five active patterns highlighted by our attribution method. The filled or hollow circles below a functional group represent whether the corresponding functional group is an active pattern for individual isozymes 1A2, 2C9, 2C19, 2D6 and 3A4 or not. The functional groups are shown on the left, while the attributed molecules with the active patterns are shown on the right.

## 5 Conclusion

In this paper, we present MoTSE, a computational method to efficiently estimate similarity between molecular property prediction tasks without any expert knowledge. Specifically, we first apply GNNs to automatically capture task-related knowledge from the corresponding datasets. Then we employ an attribution method and a molecular representation similarity analysis method to respectively extract the local and global knowledge contained in GNNs with the help of a probe dataset and project individual tasks as vectors into a latent task space. Finally, the similarity between tasks can be measured by calculating the distance between the corresponding embedded vectors in the latent task space. Comprehensive test results demonstrated that the similarity derived by MoTSE indeed can boost the performance of transfer learning on molecular property prediction, outperforming conventional training from scratch and multitask learning by a considerable margin. Moreover, by visualizing the attributed molecules, we proved that MoTSE can successfully extract the task-related knowledge and accurately estimate the intrinsic similarity between tasks, which can potentially help biologists/chemists understand the underlying biological/chemical mechanisms behind molecular properties.

## References

1. Attia, T.Z., Yamashita, T., Hammad, M.A., Hayasaki, A., Sato, T., Miyamoto, M., Yasuhara, Y., Nakamura, T., Kagawa, Y., Tsujino, H., et al.: Effect of cytochrome p450 2c19 and 2c9 amino acid residues 72 and 241 on metabolism of tricyclic antidepressant drugs. Chemical and Pharmaceutical Bulletin 62(2), 176–181 (2014)

2. Chen, G., Chen, P., Hsieh, C.Y., Lee, C.K., Liao, B., Liao, R., Liu, W., Qiu, J., Sun, Q., Tang, J., et al.: Alchemy: A quantum chemistry dataset for benchmarking ai models. arXiv preprint arXiv:1906.09427 (2019)

3. De Montellano, P.R.O.: Cytochrome P450: structure, mechanism, and biochemistry. Springer Science & Business Media (2005)

4. Dwivedi, K., Roig, G.: Representation similarity analysis for efficient task taxonomy & transfer learning. In: Proceedings of the IEEE Conference on Computer Vision and Pattern Recognition. pp. 12387–12396 (2019)

5. Eaton, E., Lane, T., et al.: Modeling transfer relationships between learning tasks for improved inductive transfer. In: Joint European Conference on Machine Learning and Knowledge Discovery in Databases. pp. 317–332. Springer (2008)

6. Fang, M., Guo, Y., Zhang, X., Li, X.: Multi-source transfer learning based on label shared subspace. Pattern Recognition Letters 51, 101–106 (2015)

7. Gaulton, A., Bellis, L.J., Bento, A.P., Chambers, J., Davies, M., Hersey, A., Light, Y., McGlinchey, S., Michalovich, D., Al-Lazikani, B., et al.: Chembl: a large-scale bioactivity database for drug discovery. Nucleic acids research 40(D1), D1100–D1107 (2012)

8. Gilmer, J., Schoenholz, S.S., Riley, P.F., Vinyals, O., Dahl, G.E.: Neural message passing for quantum chemistry. arXiv preprint arXiv:1704.01212 (2017)

9. Groen, I.I., Greene, M.R., Baldassano, C., Fei-Fei, L., Beck, D.M., Baker, C.I.: Distinct contributions of functional and deep neural network features to representational similarity of scenes in human brain and behavior. Elife 7, e32962 (2018)

10. Jain, A.K., Dubes, R.C.: Algorithms for clustering data. Prentice-Hall, Inc. (1988)

11. Jain, A.K., Murty, M.N., Flynn, P.J.: Data clustering: a review. ACM computing surveys (CSUR) 31(3), 264–323 (1999)

12. Kho, R., Hansen, M., Villar, H.: Prevalence of scaffolds in human cytochrome p450 inhibitors identified using the lopac1280 library of pharmacologically active compounds. Sigma-Aldrich URL: http://www.sigmaaldrich.com/AreaofInterest/LifeScience/LifeScienceQuarterly/Spring_2006.html (2006)

13. Kim, S., Thiessen, P.A., Bolton, E.E., Chen, J., Fu, G., Gindulyte, A., Han, L., He, J., He, S., Shoemaker, B.A., et al.: Pubchem substance and compound databases. Nucleic acids research 44(D1), D1202–D1213 (2016)

14. Kingma, D.P., Ba, J.: Adam: A method for stochastic optimization. arXiv preprint arXiv:1412.6980 (2014)

15. Kipf, T.N., Welling, M.: Semi-supervised classification with graph convolutional networks. arXiv preprint arXiv:1609.02907 (2016)

16. Kohonen, T.: Self-organized formation of topologically correct feature maps. Biological cybernetics 43(1), 59–69 (1982)

17. Kusner, M.J., Paige, B., Hernández-Lobato, J.M.: Grammar variational autoencoder. arXiv preprint arXiv:1703.01925 (2017)

18. Lee, J., Basith, S., Cui, M., Kim, B., Choi, S.: In silico prediction of multiple-category classification model for cytochrome p450 inhibitors and non-inhibitors using machine-learning method. SAR and QSAR in Environmental Research 28(10), 863–874 (2017)

19. Li, X., Yan, X., Gu, Q., Zhou, H., Wu, D., Xu, J.: Deepchemstable: chemical stability prediction with an attention-based graph convolution network. Journal of chemical information and modeling 59(3), 1044–1049 (2019)

20. Macarron, R., Banks, M.N., Bojanic, D., Burns, D.J., Cirovic, D.A., Garyantes, T., Green, D.V., Hertzberg, R.P., Janzen, W.P., Paslay, J.W., et al.: Impact of high-throughput screening in biomedical research. Nature reviews Drug discovery 10(3), 188–195 (2011)

21. Mayr, A., Klambauer, G., Unterthiner, T., Steijaert, M., Wegner, J.K., Ceulemans, H., Clevert, D.A., Hochreiter, S.: Large-scale comparison of machine learning methods for drug target prediction on chembl. Chemical science 9(24), 5441–5451 (2018)

22. Papadatos, G., Gaulton, A., Hersey, A., Overington, J.P.: Activity, assay and target data curation and quality in the chembl database. Journal of computer-aided molecular design 29(9), 885–896 (2015)

23. Ramakrishnan, R., Dral, P.O., Rupp, M., von Lilienfeld, O.A.: Quantum chemistry structures and properties of 134 kilo molecules. Scientific Data 1(2014)

24. Ramsundar, B., Kearnes, S., Riley, P., Webster, D., Konerding, D., Pande, V.: Massively multitask networks for drug discovery. arXiv preprint arXiv:1502.02072 (2015)

25. Rogers, D., Hahn, M.: Extended-connectivity fingerprints. Journal of chemical information and modeling 50(5), 742–754 (2010)

26. Rosenstein, M.T., Marx, Z., Kaelbling, L.P., Dietterich, T.G.: To transfer or not to transfer. In: NIPS 2005 workshop on transfer learning. vol. 898, pp. 1–4 (2005)

27. Schneider, P., Schneider, G.: Collection of bioactive reference compounds for focused library design. QSAR & Combinatorial Science 22(7), 713–718 (2003)

28. Selzer, P., Ertl, P.: Applications of self-organizing neural networks in virtual screening and diversity selection. Journal of chemical information and modeling 46(6), 2319–2323 (2006)

29. Shen, J., Nicolaou, C.A.: Molecular property prediction: recent trends in the era of artificial intelligence. Drug Discovery Today: Technologies (2020)

30. Shrikumar, A., Greenside, P., Shcherbina, A., Kundaje, A.: Not just a black box: Learning important features through propagating activation differences. arXiv preprint arXiv:1605.01713 (2016)

31. Simoes, R.S., Maltarollo, V.G., Oliveira, P.R., Honorio, K.M.: Transfer and multi-task learning in qsar modeling: advances and challenges. Frontiers in pharmacology 9, 74 (2018)

32. Sterling, T., Irwin, J.J.: Zinc 15–ligand discovery for everyone. Journal of chemical information and modeling 55(11), 2324–2337 (2015)

33. Veith, H., Southall, N., Huang, R., James, T., Fayne, D., Artemenko, N., Shen, M., Inglese, J., Austin, C.P., Lloyd, D.G., et al.: Comprehensive characterization of cytochrome p450 isozyme selectivity across chemical libraries. Nature biotechnology 27(11), 1050–1055 (2009)

34. Veličković, P., Cucurull, G., Casanova, A., Romero, A., Lio, P., Bengio, Y.: Graph attention networks. arXiv preprint arXiv:1710.10903 (2017)

35. Wang, M., Zheng, D., Ye, Z., Gan, Q., Li, M., Song, X., Zhou, J., Ma, C., Yu, L., Gai, Y., Xiao, T., He, T., Karypis, G., Li, J., Zhang, Z.: Deep graph library: A graph-centric, highly-performant package for graph neural networks. arXiv preprint arXiv:1909.01315 (2019)

36. Weininger, D.: Smiles, a chemical language and information system. 1. introduction to methodology and encoding rules. Journal of chemical information and computer sciences 28(1), 31–36 (1988)

37. Williams, J.A., Hyland, R., Jones, B.C., Smith, D.A., Hurst, S., Goosen, T.C., Peterkin, V., Koup, J.R., Ball, S.E.: Drug-drug interactions for udp-glucuronosyltransferase substrates: a pharmacokinetic explanation for typically observed low exposure (auci/auc) ratios. Drug Metabolism and Disposition 32(11), 1201–1208 (2004)

38. Xiong, Z., Wang, D., Liu, X., Zhong, F., Wan, X., Li, X., Li, Z., Luo, X., Chen, K., Jiang, H., et al.: Pushing the boundaries of molecular representation for drug discovery with the graph attention mechanism. Journal of Medicinal Chemistry (2019)

39. Zamir, A.R., Sax, A., Shen, W., Guibas, L.J., Malik, J., Savarese, S.: Taskonomy: Disentangling task transfer learning. In: Proceedings of the IEEE conference on computer vision and pattern recognition. pp. 3712–3722 (2018)

40. Zhang, W., Deng, L., Wu, D.: Overcoming negative transfer: A survey. arXiv preprint arXiv:2009.00909 (2020)

