## Supplementary Notes and Figures for "MoTSE: an interpretable task similarity estimator for small molecular property prediction tasks"

### 1 Supplementary Details on Experimental Settings

#### 1.1 Datasets

| Dataset | Tasks |
|---|---|
| QM9 | mu, alpha, homo, lumo, gap, r2, zpve, u0, u298, h298, g298, cv |
| PCBA | PCBA-1030, PCBA-1458, PCBA-1460, PCBA-2546, PCBA-2551, PCBA-485297, PCBA-485313, PCBA-485364, PCBA-504332, PCBA-504333, PCBA-504339, PCBA-504444, PCBA-504467, PCBA-588342, PCBA-624296, PCBA-624297, PCBA-624417, PCBA-651965, PCBA-652104, PCBA-686970, PCBA-686978, PCBA-686979, PCBA-720504 |

**Table S1.** Tasks in the preprocessed QM9 and PCBA datasets. The preprocessed QM9 dataset contains 12 tasks measuring quantum chemistry properties of molecules. The preprocessed PCBA dataset contains 23 tasks measuring biological activities of molecules.

In Section 3 of the main text, we mainly used two datasets, including QM9 and PCBA, to evaluate the performance of MoTSE on estimating task similarity. The tasks associated with these two datasets are shown in Table S1. In addition to the datasets created in Section 3.1, to further validate the robustness of MoTSE, we also created a set of imbalanced PCBA dataset containing 10k samples, denoted as $\text{PCBA}_{p\_10k}$, where $p \in \{10\%, 20\%, 30\%, 40\%\}$ stands for the proportion of positive samples in the dataset. For each $p \in \{10\%, 20\%, 30\%, 40\%\}$, we constructed $\text{PCBA}_{p\_10k}$ by randomly sampling $p \times 10k$ positive and $(1-p) \times 10k$ negative samples for individual tasks in the preprocessed PCBA data as shown in Table S1. Then these datasets were partitioned into training (8k), validation (1k) and test (1k) sets. Correspondingly, for each $p \in \{10\%, 20\%, 30\%, 40\%\}$, we also created $\text{PCBA}_{p\_n}$ by randomly sampling $0.8 \times n$ and $0.1 \times n$ samples from the training and validation sets of $\text{PCBA}_{p\_10k}$, respectively, where $n \in \{1k, 2k, 3k, 4k\}$ stands for the size of the dataset. The testing set of $\text{PCBA}_{p\_n}$ was the same as that of $\text{PCBA}_{p\_10k}$ for a fair comparison. During transfer learning, we used $\text{PCBA}_{p\_10k}$ for pretraining and $\text{PCBA}_{p\_n}$ for finetuning.

#### 1.2 Molecular Representations

In addition to molecular graph, we also used ECFP (extended connectivity fingerprints) [7] and SMILES (simplified molecular input line entry specification) [9] to represent molecules in our tests. ECFP is a

⋆ This work was supported in part by the National Natural Science Foundation of China [61872216, 81630103], the Turing AI Institute of Nanjing and the Zhongguancun Haihua Institute for Frontier Information Technology.

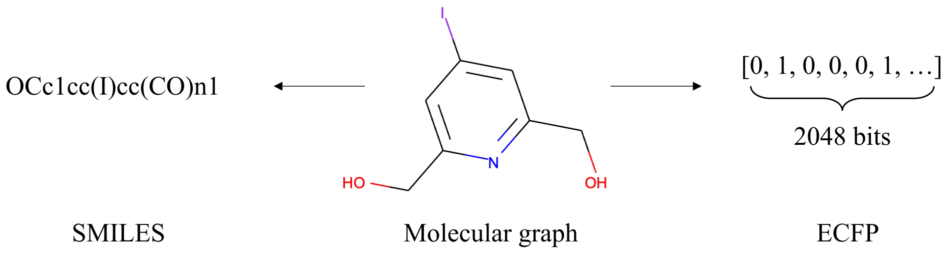


**Figure S1.** Three types of feature representations of an example molecule. ECFP denotes extended connectivity fingerprints. SMILES denotes simplified molecular input line entry specification.

representation of 2D binary fingerprints (i.e., a series of bits) which can dynamically index the presence or absence of particular substructures in molecules. SMILSE is a string-based molecular representation for describing molecular structures using short ASCII strings. Figure S1 gives an example for both ECFP and SMILES representations. In our experiments, we used DeepChem [6] to calculate a 2048-bit ECFP for each molecule. For SMILES string, we used a one-hot vector to encode the unique characters.

##### 1.3 Models

To evaluate whether the similarity estimated using a GCN can be generalized to guide the source task selection of other models, we also employed a graph attention network (GAT) [8], a fully-connected network (FCN) and a recurrent neural network (RNN). The details of these models are provided below.

- GCN: We constructed the GCN model with three 256-dimensional GCN layers followed by a two-layer (256-128-1) fully connected network for molecular representation extraction and property prediction.
- GAT: GAT is another kind of graph neural network which employs attention mechanism when performing message passing over nodes. We constructed the GAT model with three 256-dimensional GAT layers followed by a two-layer (256-128-1) fully connected network for molecular representation extraction and prediction.
- FCN: We built a five-layer (2048-1024-512-256-128-1) fully connected network which takes the ECFP of molecules as input to make molecular property prediction.
- RNN: RNN is a deep learning model particularly designed for processing sequential data, which has been proven to be effective in making molecular property predictions with one-hot encoded SMILES [1, 2]. Here we employed a three-layer 128-dimensional LSTM [3] (a classical variant of RNN) to encode SMILES representations of molecules to 64-dimensional latent vectors and a two-layer (64-32-1) fully connected neural network to make prediction.

##### 1.4 Pretraining and Finetuning

During pretraining and finetuning, we preserved the same hyper-parameters across all tasks and models. All the models was trained with the Pytorch [5] framework. MSELoss and CrossEntropyLoss were employed to measure the mean-squared error and the cross entropy for the regression tasks and classification tasks, respectively. We used Adam optimizer [4] for gradient descent optimization with the following hyper-parameters: learning rate is set to 1e-4 and weight decay is set to 1e-5. All models were trained for 1000 epochs with early stopping which aims to terminate training when the validation accuracy has not been improved in the last 20 epochs.

#### 2 Supplementary Experimental Results

##### 2.1 Supplementary Details on Performance Evaluation

In this section, we showed that by increasing the number of tasks selected in the source task selection process, we can gradually improve the prediction performance of transfer learning with the guidance of

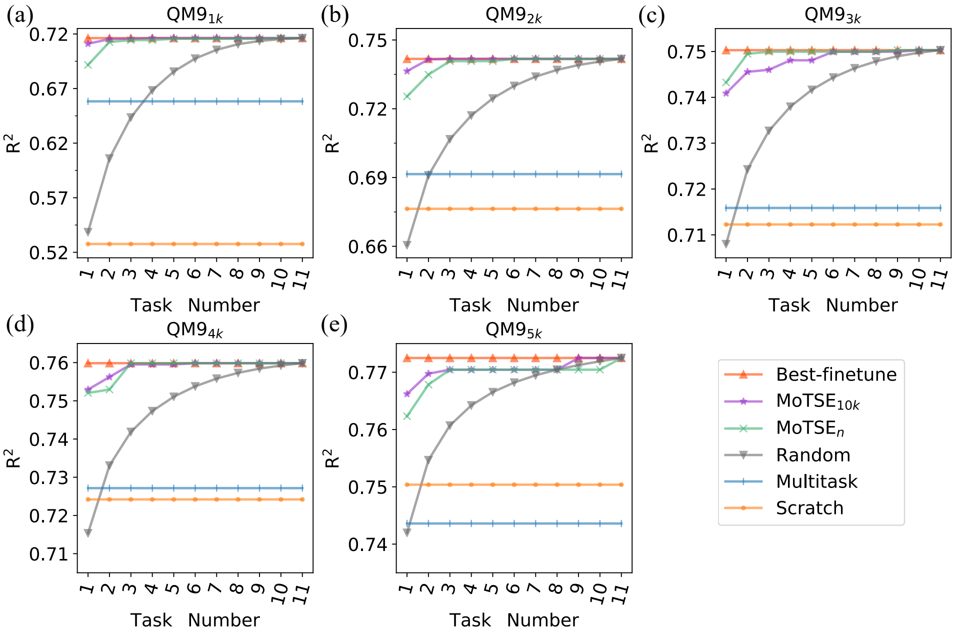


**Figure S2.** Improving the prediction performance of transfer learning on QM9$_n$ by selecting more tasks according to MoTSE$_{10k}$ and MoTSE$_n$, where $n \in \{1k, 2k, 3k, 4k\}$.

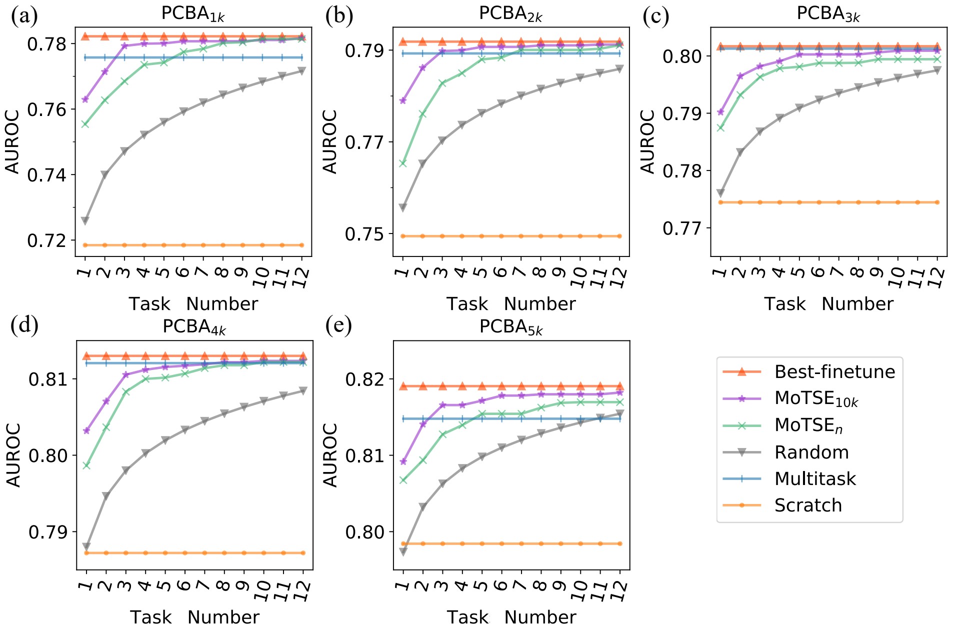


**Figure S3.** Improving the prediction performance of transfer learning on PCBA$_n$ by selecting more tasks according to MoTSE$_{10k}$ and MoTSE$_n$, where $n \in \{1k, 2k, 3k, 4k\}$.

similarity estimated by MoTSE (i.e., $\text{MoTSE}_{10k}$ and $\text{MoTSE}_n$) until the best performance (i.e., Best-finetune) was reached. Using $\text{QM9}_{10k}$ and $\text{PCBA}_{10k}$ for pretraining, for each $n \in \{1k, 2k, 3k, 4k, 5k\}$, we plotted the number of selected source tasks versus $R^2$ and AUROC derived from finetuning on $\text{QM9}_n$ (Figure S2) and $\text{PCBA}_n$ (Figure S3). We also provided the results of Best-finetune, Random, Multitask, Scratch as defined in Section 3.2 for individual tests.

##### 2.2 Supplementary Details on Evaluating the Robustness of MoTSE

In Section 3.2, we have shown that the similarity estimated by MoTSE using limited data can also effectively guide the source task selection (Figure 2), which demonstrated that MoTSE is robust even given limited data. In this section, we further evaluate whether MoTSE is robust given different probe datasets or imbalanced datasets.

Probe dataset is an importance part in our MoTSE framework, which was shared across all tasks and acted as a proxy in the process of projecting individual tasks into the latent task space as described in Section 2. To evaluate the robustness of MoTSE when given different probe datasets, we first randomly sampled three probe datasets from the preprocessed Zinc dataset and used MoTSE to estimate similarity between tasks in $\text{QM9}_{10k}$ and $\text{PCBA}_{10k}$ using these three probe datasets, respectively. Then we measured the Pearson's and Spearman's correlations between the similarity estimated using three different probe datasets for individual tasks. The average of Pearson's and Spearman's correlations across all tasks on the QM9 were 0.999 and 0.977, respectively. The average of Pearson's and Spearman's correlations across all tasks on PCBA were 0.996 and 0.886, respectively. Such high correlation results indicated that MoTSE is robust when given different probe datasets.

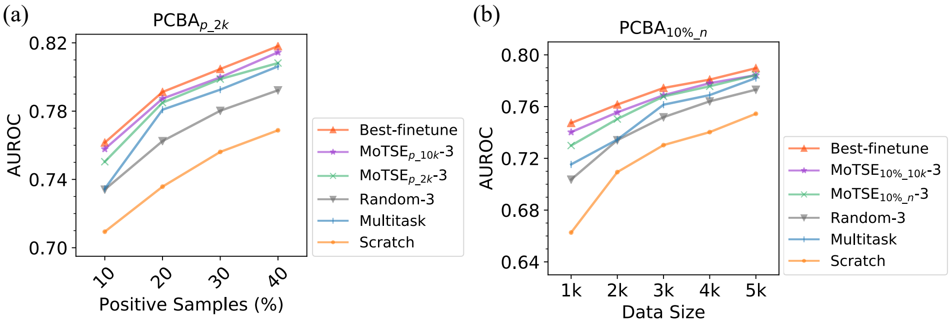


**Figure S4.** Prediction performance on imbalanced dataset of transfer learning with the guidance of the similarity estimated by MoTSE and other strategies. (a) The prediction performance on $\text{PCBA}_{p\_2k}$, where $p \in \{10\%, 20\%, 30\%, 40\%\}$ stands for the proportion of the positive data in dataset. (b) The prediction performance on $\text{PCBA}_{10\%\_n}$, where $n \in \{1k, 2k, 3k, 4k\}$ is the size of the dataset for finetuning.

To evaluate the robustness of MoTSE when given imbalanced datasets, we also carried out the following two additional tests. (1) For each $p \in \{10\%, 20\%, 30\%, 40\%\}$, we used MoTSE to estimate task similarity using $\text{PCBA}_{p\_10k}$ and $\text{PCBA}_{p\_2k}$, denoted as $\text{MoTSE}_{p\_10k}$ and $\text{MoTSE}_{p\_2k}$, respectively. Then $\text{MoTSE}_{p\_10k}$ and $\text{MoTSE}_{p\_2k}$ were used to guide the transfer learning process when using $\text{PCBA}_{p\_10k}$ for pretraining and $\text{PCBA}_{p\_2k}$ for finetuning. (2) For each $n \in \{1k, 2k, 3k, 4k\}$, we used MoTSE to estimate task similarity using $\text{PCBA}_{10\%\_10k}$ and $\text{PCBA}_{10\%\_n}$, denoted as $\text{MoTSE}_{10\%\_10k}$ and $\text{MoTSE}_{10\%\_n}$, respectively. Then $\text{MoTSE}_{10\%\_10k}$ and $\text{MoTSE}_{10\%\_n}$ were used to guide the transfer learning process when using $\text{PCBA}_{10\%\_10k}$ for pretraininig and $\text{PCBA}_{10\%\_n}$ for finetuning. For the first experiment, we plotted the proportion ($p$) of positive samples versus AUROC on $\text{PCBA}_{p\_2k}$ (Figure S4a). For the second experiment, we plotted the data size ($n$) versus AUROC on $\text{PCBA}_{10\%\_n}$ (Figure S4b). We also provided the results of Best-finetune, Random-3, Scratch and Multitask as defined in Section 3.2 for individual tests.

#### 2.3 Supplementary Information on Analyzing the Efficiency of MoTSE

In Section 3.1, we briefly explained that MoTSE estimates task similarity in a more efficient manner than performing transfer learning for all pairs of tasks (i.e., estimating task transferability). Here, we gave a more mathematically analysis on the computational complexity of these two methods.

Given a task set $\mathcal{T}$ with $N_{\mathcal{T}}$ tasks and corresponding datasets $\mathcal{D}$ with $N_{\mathcal{D}}$ data size for each dataset $D \in \mathcal{D}$, assume that the prediction model is finetuned for $E$ epochs. Note that, we omitted the pretraining process because it performs in both methods. We also excluded the cost of similarity estimation as it can be finished within 10 minutes in our experiments. Then the forward-and-backward propagation (FPP) times of estimating task transferability can be approximately formulized as:

$$T_{Transfer} = N_{\mathcal{T}}(N_{\mathcal{T}} - 1)N_{\mathcal{D}}E. \tag{1}$$

For MoTSE, it only requires one time FPP on the probe dataset. Assume that the size of probe dataset is $N_p$. Then the FPP times of MoTSE is:

$$T_{MoTSE} = N_{\mathcal{T}}N_p. \tag{2}$$

Therefore, MoTSE is about $\frac{(N_{\mathcal{T}}-1)N_{\mathcal{D}}E}{N_p}$ times more efficient than task transferability estimation, which is a significant improvement since usually $N_p < N_{\mathcal{D}}$ and $N_{\mathcal{T}}$ can be quite large.

#### 2.4 Supplementary Details on the Constructed Similarity Tree

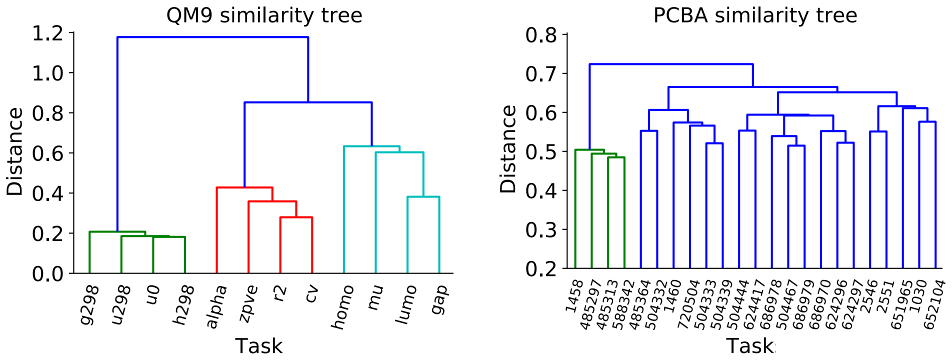


**Figure S5.** The similarity tree of individual tasks in QM9 and PCBA.

We constructed similarity trees for QM9 and PCBA tasks using agglomerative hierarchical clustering based on the similarity estimated by MoTSE. As shown in Figure S5, MoTSE explicitly clustered tasks in QM9 into three groups with low intra-class distance and high inter-class distance, while the groups of PCBA tasks were not as clear as in the QM9 case. This may be explained by the observation that the intrinsic similarity between the tasks in QM9 is relatively explicit. For example, the tasks including g298, u298, u0 and h298 in the QM9 dataset all measure the thermodynamic properties of molecules. However, the similarity between biological activity tasks (i.e., the tasks in PCBA) is much more complicated mainly due to the high complexity of the human body system.

#### 2.5 Effects of Hyper-Parameters

**Effect of the Value of $\boldsymbol{\lambda}$** As described in Section 2.3, We use $\lambda$ to adjust the weights of the extracted local knowledge and global knowledge in the similarity estimation. In this section, we evaluated the influence of the value of $\lambda$ on the model performance. More specifically, we used $\text{PCBA}_{10k}$ and $\text{QM9}_{10k}$

for pretraining, while using $PCBA_{1k}$ and $QM9_{1k}$ for finetuning. Then we plotted the values of $\lambda$ versus AUROC and $R^2$ on $PCBA_{1k}$ and $QM9_{1k}$ (Figure S6a). The results showed that the PCBA dataset preferred a larger $\lambda$, while QM9 dataset preferred a smaller $\lambda$. This observation was consistent with the conclusion that we draw in Section 3.2 (i.e., bioacitivity properties in PCBA are mostly determined by the local features, e.g., functional groups, while some of the properties in QM9 are mostly determined by the global features).

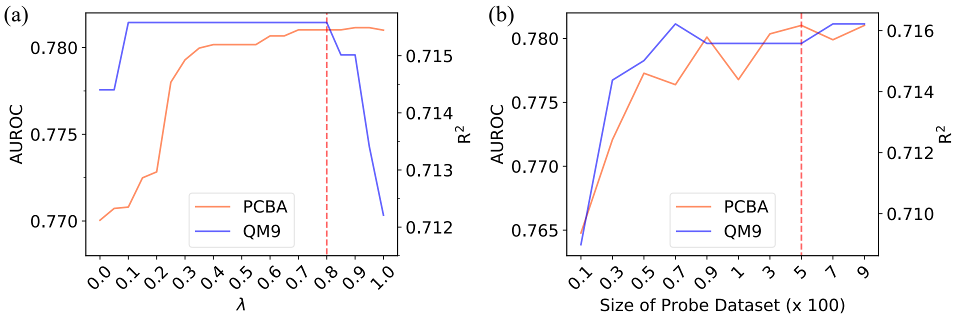


**Figure S6.** The transfer learning performance on $PCBA_{1k}$ and $QM9_{1k}$ measured in terms of AUROC and $R^2$, when recommending source tasks using $MoTSE_{10k}$ with different values of $\lambda$ (a) and different sizes of the probe dataset (b), respectively.

**Effect of the Size of Probe Dataset** In this section, we investigated the effect of the size of probe dataset on the recommendation performance. Using $PCBA_{10k}$ and $QM9_{10k}$ for pretraining, we showed the transfer learning performance on $PCBA_{1k}$ and $QM9_{1k}$ when recommending source tasks using $MoTSE_{10k}$ with different sizes of probe dataset (Figure S6b). From the results, the recommendation performance kept stable at the high accuracy when the size of probe dataset larger than 300.

### References

1. Arús-Pous, J., Johansson, S.V., Prykhodko, O., Bjerrum, E.J., Tyrchan, C., Reymond, J.L., Chen, H., Engkvist, O.: Randomized smiles strings improve the quality of molecular generative models. Journal of cheminformatics **11**(1), 1–13 (2019)
2. Goh, G.B., Hodas, N., Siegel, C., Vishnu, A.: Smiles2vec: Predicting chemical properties from text representations (2018)
3. Hochreiter, S., Schmidhuber, J.: Long short-term memory. Neural computation **9**(8), 1735–1780 (1997)
4. Kingma, D.P., Ba, J.: Adam: A method for stochastic optimization. arXiv preprint arXiv:1412.6980 (2014)
5. Paszke, A., Gross, S., Chintala, S., Chanan, G., Yang, E., DeVito, Z., Lin, Z., Desmaison, A., Antiga, L., Lerer, A.: Automatic differentiation in pytorch (2017)
6. Ramsundar, B., Eastman, P., Walters, P., Pande, V., Leswing, K., Wu, Z.: Deep Learning for the Life Sciences. O'Reilly Media (2019), https://www.amazon.com/Deep-Learning-Life-Sciences-Microscopy/dp/1492039837
7. Rogers, D., Hahn, M.: Extended-connectivity fingerprints. Journal of chemical information and modeling **50**(5), 742–754 (2010)
8. Veličković, P., Cucurull, G., Casanova, A., Romero, A., Lio, P., Bengio, Y.: Graph attention networks. arXiv preprint arXiv:1710.10903 (2017)
9. Weininger, D.: Smiles, a chemical language and information system. 1. introduction to methodology and encoding rules. Journal of chemical information and computer sciences **28**(1), 31–36 (1988)
